# Experience, not time, determines representational drift in the hippocampus

**DOI:** 10.1101/2022.08.31.506041

**Authors:** Dorgham Khatib, Aviv Ratzon, Mariell Sellevoll, Omri Barak, Genela Morris, Dori Derdikman

## Abstract

Memories of past events can be recalled long after the event, indicating stability. But new experiences are also integrated into existing memories, indicating plasticity. In the hippocampus, spatial representations are known to remain stable, but have also been shown to drift over long periods of time. We hypothesized that experience, more than the passage of time, is the driving force behind memory plasticity. We compared the stability of place cells in the hippocampus of mice traversing two similar, familiar tracks for different durations. We found that the more time spent in an environment, the greater the representational drift, regardless of the total elapsed time. Our results suggest that spatial representation is a dynamic process, related to the ongoing experiences within a specific context, and is related to the accumulation of new memories rather than to passive forgetting.

**Highlights:**

- Representational drift is related to experience within an environment.
- Representational drift is a dynamic context-wide process.
- Place cell number decreases with experience, spatial information content increases.

## INTRODUCTION

Place cells in the hippocampus^1–3^ are thought to be involved in the representation of episodic memories^4,5^. When recalling a memory that involves the hippocampus, the memory is reinstated in the pattern of cell activity in the CA1, according to the synaptic strengths at the moment of encoding^6,7^. To represent such memories, place cells must be both stable enough to hold the core memory^8^, yet dynamic enough to allow the introduction of changes to the memory, thus enabling memory updating^9^.

Indeed, the representation of place cells in the hippocampus has been shown to gradually change over time, within the same context. Referred to as gradual remapping^8,10,11^, and also as representational drift, this process occurs over a period of hours to days, of repeated exposures to the same environment^12–15^, although it has been shown that context representation may be preserved^16^. Two mechanisms could potentially contribute to representational drift. One mechanism is time dependent, whereby the passage of time weakens memories, leading to partial forgetting of the original representation. The second mechanism involves context updating, by which memories that are more malleable to change, due to the specific context and the amount of experience accumulated within them, are continuously updated. The question arises as to whether the malleability of memories is more affected by their relative use, or rather by the absolute passage of time.

To address this question, we used a behavioral paradigm that dissociates time and experience. A mouse traversed two familiar connected environments in the same imaging session. The mouse visited one of the environments for only short periods at the beginning and at the end of the session, while spending the remaining hours of the session in the other environment. Thus, the absolute time interval between the first and last recording in each environment was identical, while the time spent in each environment differed by an order of magnitude. We found that spatial representations in the hippocampus changed differently in the two environments, such that the rate of change depended on the level of use: the longer a memory in a certain context was active, the more malleable it was to change.

## RESULTS

### Context-dependent change of representation is a function of accumulated experience in CA1

We set out to distinguish between the effects of passage of time and experience in a context, on the representational drift in the hippocampus. Thus, we designed a U-Shaped Maze consisting of two linear tracks of similar length, connected by an intermediate chamber with two doors that open to either track (Figure 1A). Mice were trained to traverse both tracks, running back and forth, and collecting food rewards at both ends of the track. The habituation period was 4-6 days; on each day the animals ran 20 minutes in each arm of the maze until they were consistently able to run back and forth to collect the food rewards.

**Figure 1.**
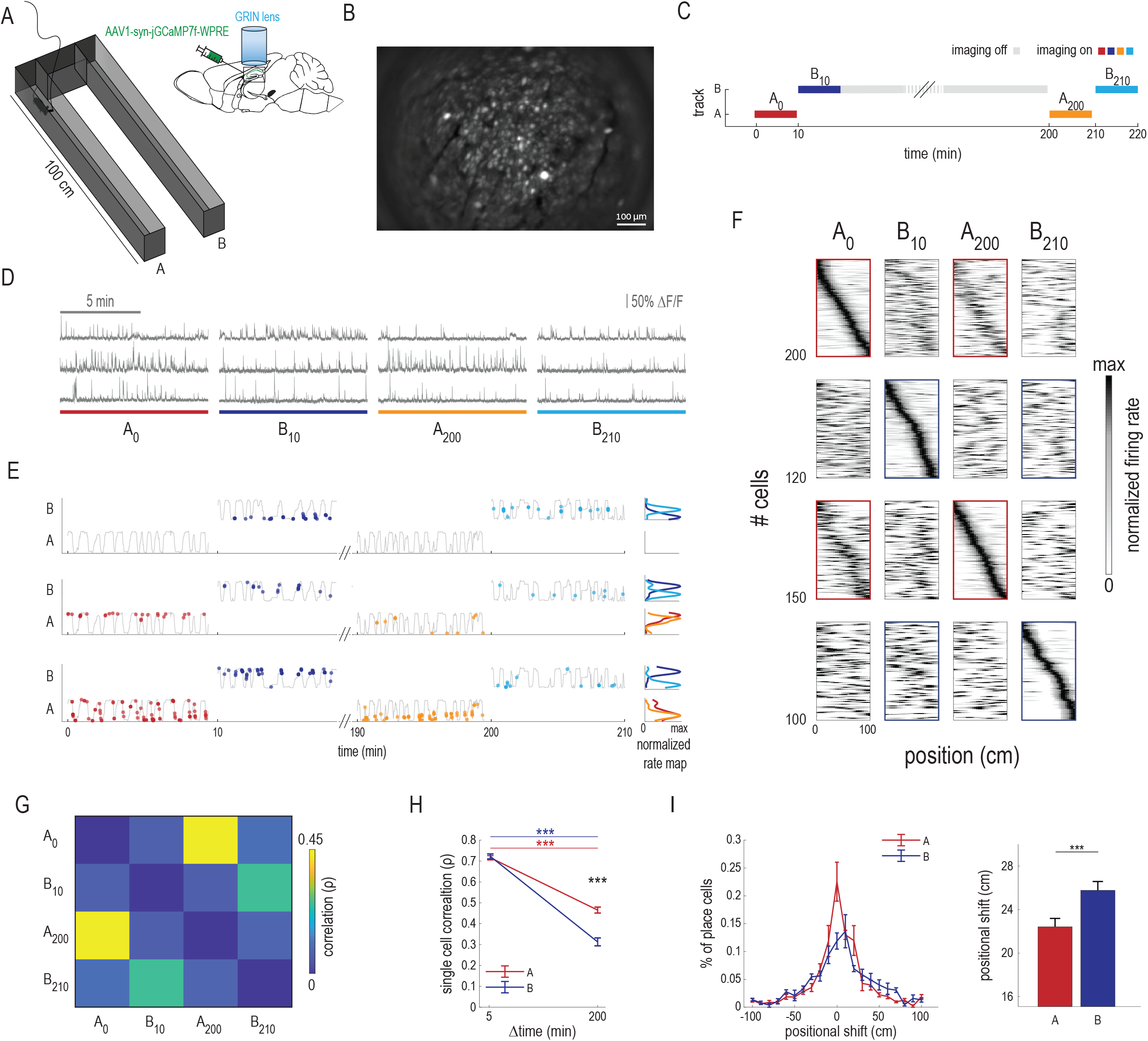
Context dependent change of representation as a function of the relative time in the dorsal CA1. A. Left: Illustration of the maze, composed of two linear tracks, A and B, connected by an intermediate chamber. Right: Illustration of the injection site and viral vector used, together with the lens implantation. B. Example of an imaging field of view. C. The timeline of the experiment. The colors correspond to the different recordings; grey color indicates experiment time without imaging. D. Three examples of place cell calcium traces in all the recordings. E. Examples of three place cells, showing the firing rate of the cells as a function of position in the track (right) and the Ca^2+^ events (left) along the different recordings (color code as in B). F. Place cell population in all the recordings for mouse 9855. In each row are shown average rate maps for individual cells along the linear track, during left or right traversals of place cells that were active in that specific period, normalized by peak activity. In the n^th^ row, place cells were selected and sorted according to the n^th^ imaging session. Note that each column displays data from the same session. The red and blue rectangles emphasize the comparison between the first and last sessions in tracks A and B, respectively. G. Pairwise Spearman correlations of the individual place cells shown in “e”, between recordings. H. Pairwise correlations of all the place cells from all the mice (n=3 mice, 1441 cells) as a function of elapsed time. The 5-minute point was computed by splitting the recordings in half and correlating the activity of place cells between consecutive halves. The correlations in track B were lower than in track A (mean±s.e.m: A_200_=0.46±0.01; B_210_=0.31±0.01; p<0.001, Kruskal-Wallis with Dunn’s test). I. Left: Positional shift of the center of mass of the place cells of all the mice (n=3, 1441 cells), between the first and last recordings in tracks A and B. Right: The overall positional shift of the center of mass of the place cells, between tracks A and B. Positional shift increased in track B compared to track A (mean±s.e.m: A_200_=22.46±0.76; B_210_=25.77±0.8; p<0.001 Mann-Whitney U test).

We performed Ca^2+^ imaging of neuronal activity in the dorsal CA1 (dCA1) in freely moving mice using a one-photon miniature microscope and a micro-endoscope probe (*mean* ± *SD:* 456 ± 146 cells per session) (Figure 1B). On the day of the experiment, the mice were first placed inside the intermediate chamber. Subsequently, they were released into track A, where they spent 10 minutes, while their neuronal activity was imaged (A_0_). We then opened the doors into track B, which the mice traversed for 200 minutes. The activity of cells in track B was imaged for the first 10 minutes only (B_10_). Subsequently, the mice returned to track A through the intermediate chamber for 10 additional minutes of imaging (A_200_), and finally to track B where the cells were imaged for an additional 10 minutes (B_210_). This yielded a total of four 10-minute recordings (Figure 1C). The experimental design ensured that the absolute time that had passed between A_0_ and A_200_ was identical to the absolute time passed between B_10_ and B_210_.

Visually comparing activity (Figure 1D, E) across recordings indicated greater resemblance in the spatial maps of track A (A_0_ and A_200_) than of track B (B_10_ and B_210_) (Figure 1F, Supplementary figure 1A). To quantify changes over time in the representations in tracks A and B, we calculated the correlations between the population rate maps in each pair of recordings (Figure 1G, Supplementary figure 1B). To obtain a baseline correlation value, within a recording period, we divided each 10-minute period into two five-minute periods and correlated these with each other. We found a significantly lower correlation between the maps in the B_10_ and B_210_ recordings than in the maps of the A_0_ and A_200_ recordings (Figure 1H, Supplementary figure 1C). To quantify the extent of representational drift, we calculated the positional shift of the center of mass of each place field, between the first and last recordings in each track. This yielded a significant increase in positional shift of place cells in track B, compared to track A (Figure 1I, Supplementary figure 1D). Overall, these results indicate greater change in representation of a context while the animal is in that context, relative to when it is not.

### Representational drift is a gradual process

To investigate the dynamics of the representational drift observed in track B, we repeated the behavioral protocol described above, while introducing repetitive recordings. Specifically, we performed 10-minute recordings every 50 minutes in track B, resulting in a total of five recordings (B_10_, B_70_, B_130_, B_190_ and B_210_), and two recordings in track A (A_0_ and A_200_) (Figure 2A). Rate maps of place cells underwent gradual change, from B_10_ to B_210_ (Figure 2B). This was also reflected in a gradual decrease in inter-map correlations as a function of the time interval between these maps (Figure 2C, D, Supplementary figure 2A, B). In this experiment, we reproduced the initial finding, namely, higher correlations between the first and last recordings in track A compared to the first and last recordings in track B (Figure 2E, Supplementary figure 2C). The positional shift increased as the correlations decreased (Figure 2F, G, Supplementary figure 2D).

**Figure 2.**
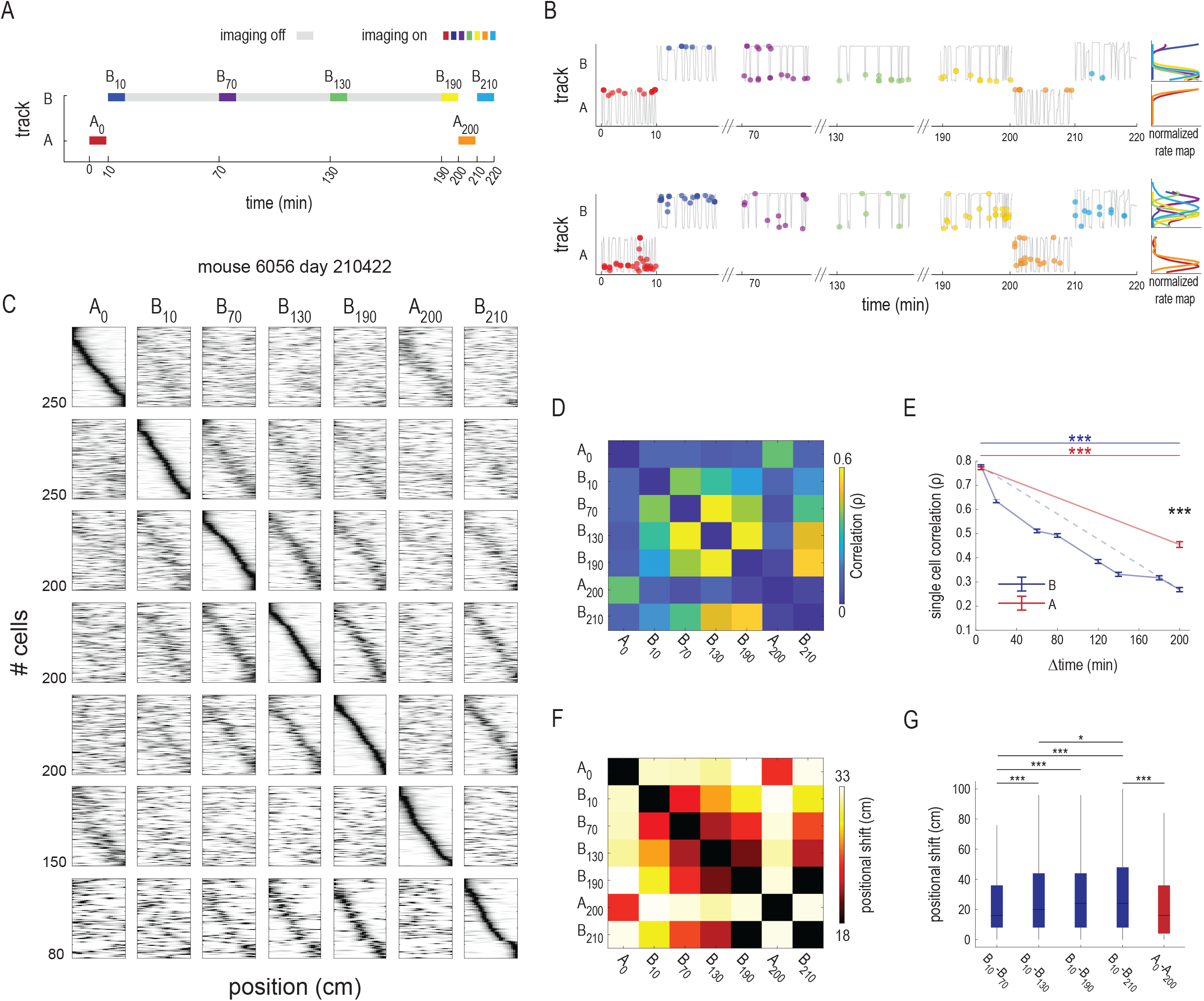
Representational drift is a gradual process. A. Timeline of the experiment, the colors correspond to the different recordings, grey color indicates experiment time with imaging off. B. Examples of two place cells, showing the firing rate of the cells as a function of their positions in the track (right) and the Ca^2+^ event times as a function of position (left), over the course of the recordings (same colors as in panel a). C. The place cell population in all the imaging sessions for mouse 6065. The maps show average rate maps for individual cells along the right or left traversals of the linear track of the place cells that were active during the specific recording, normalized by the peak activity. In the n^th^ row, place cells are selected and sorted according to the n^th^ recording. Note that each column displays data from the same recording. D. Average pairwise correlations of individual cells, between imaging sessions for the same mouse. E. Pairwise correlation of all place cells (n=5 mice, 4280 cells) as a function of elapsed time. Correlations in track A were higher than in track B (mean±s.e.m: A_200_=0.45±0.01; B_210_=0.26±0.008; p<0.001, Kruskal-Wallis with Dunn’s test). F. An example of positional shift of the center of mass of all place cells from the same mouse shown in “c”. G. Positional shift compared between recordings in track A and recordings in track B, p<0.001, Kruskal-Wallis with Dunn’s test.

### Representational drift is a context-wide process

We were interested in further investigating the extent of the context-related representational drift and whether it is related to specific sub-contexts within the bigger one. To check this, we needed a more variable behavior and a different occupancy time than is possible in a linear track. Thus, we repeated the experiment in two rectangular arenas connected by a door (Figure 3A). We used the same extended experimental protocol as before (Figure 2A). We recorded the activity in dCA1 in the two dimensional (2D) shaped arenas (Figure 3B), and we observed a similar effect. The correlations of rate maps were significantly greater for place cells in context A than for place cells in context B, albeit with a smaller effect size than in the linear track (Figure 3C, D).

**Figure 3.**
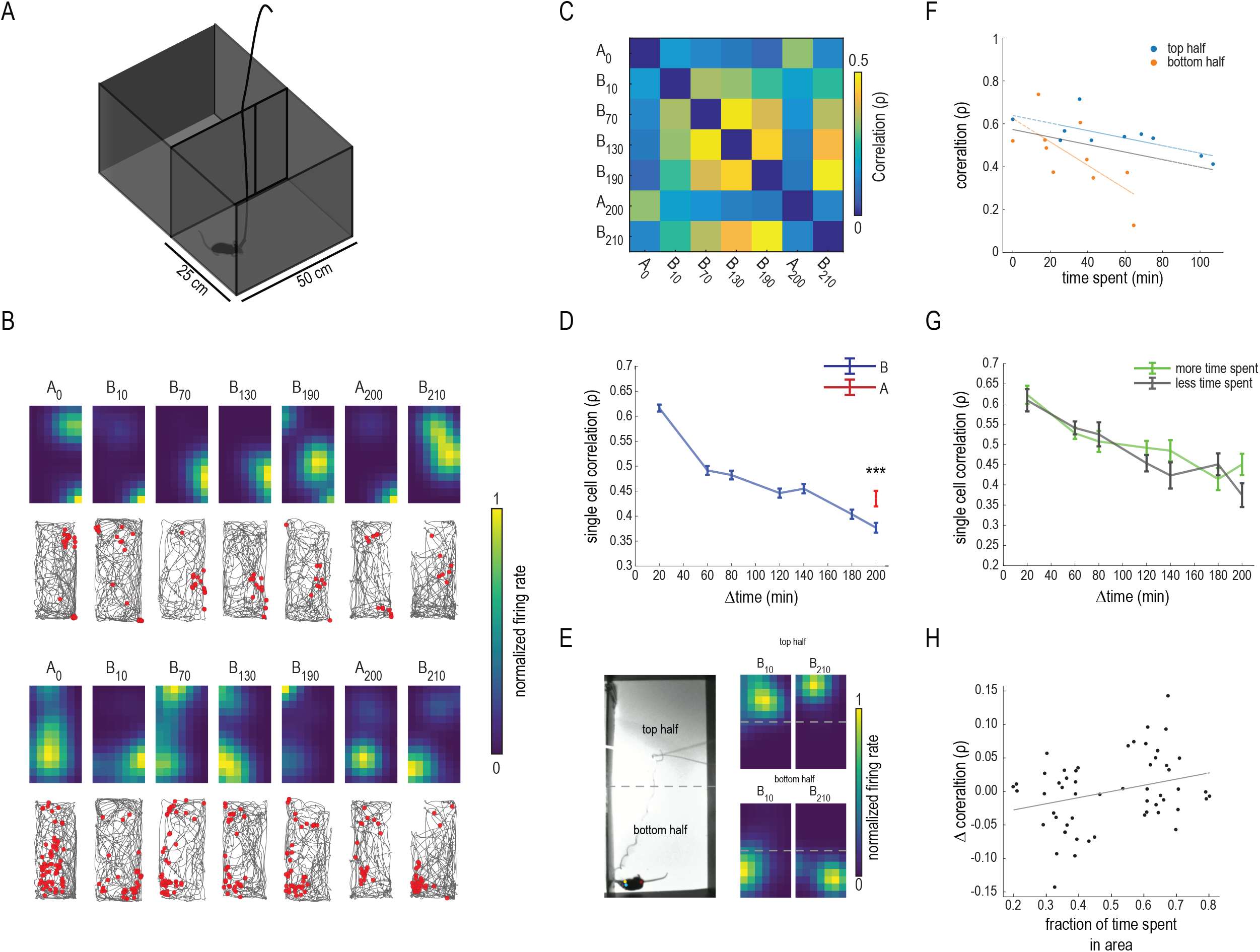
Representational drift is a context-wide process. A. Illustration of the maze. B. Two examples of place cells from all the recordings. In each example, top row: rate maps as a function of position in the track; bottom row: trajectory of the animal (black) and Ca^2+^ events (red). C. Pairwise single-cell correlations between imaging sessions, in both tracks, for mouse 9819. D. Pairwise correlation of all the place cells from all the mice (n=3 mice, 1829 cells) as a function of the time passed. Correlations after 200 min. were lower in track B than track A (mean±s.e.m: A_200_=0.40±0.01; B_210_=0.35 ±0.01, p<0.001; Kruskal-Wallis with Dunn’s test). E. Left: Illustration of the division of the arena into halves. Right: Two examples of place cells in the first and last imaging sessions in track B. F. Correlation from the top and bottom halves of the arena, for mouse 9819. The top half slope=-0.002, p=0.019; the bottom half slope=-0.005, p=0.026, two tailed t-test. G. Pairwise correlations of all the place cells from all the mice (n=3 mice, 1676 cells), as a function of the time passed. The two lines correspond to neurons whose maximal firing rate occurred in the sub-region where the mouse spent more time (green) and less time (gray). There was no significant difference between the two halves (p=0.075, Kruskal-Wallis with Dunn’s test). H. Normalized correlations as a function of the normalized time spent in each half of the arena, for all (n=3) mice. Slope=0.117, p=0.081, two tailed t-test.

Our results thus far indicate a role of experience within a specific context, in accelerating representational drift. We sought to examine whether this was also true for sub-regions. To examine this, we divided the 2D arena in arena B to two virtual halves and checked the representational drift in each half. For this analysis, we focused on the subset of neurons whose maximum firing field was initially in each sub-region (see the example in Figure 3E). We noticed that in many instances the reduction in correlation was not related to the time spent in each sub-region (e.g. Figure 3F for mouse 9819). We grouped the neurons by each mouse’s preferred sub-region, i.e. the region in which it spent more time, and measured the correlations between the rate maps of these neurons, for each pair of recordings. No significant dependence was found between the time spent in a sub-region and the reduction in correlation (Figure 3G). There was even a slight non-significant tendency for an increase in correlation when the time spent in a sub-region increased (Figure 3H). Note that in the specific experimental setting, this analysis cannot be reliably done in a linear track, as the mice tended to spend an equal amount of time throughout such a track. In the 2D arena, each mouse had a preferred sub-region where it spent more time, thus enabling testing the hypothesis. In summary, representational drift appears to occur as a single entity in the entire context, unrelated to the occupancy time within sub-regions of the context.

### Spatial information content of place cells increases while the number of place cells decreases, as more time is spent in an environment

To understand how representational drift affects the spatial encoding of the environment, we measured parameters of spatial information and encoding. We examined the spatial information content of place cells in each of the recordings. Spatial information of place cells increased with the time spent in the environment (Figure 4A left). The increase in spatial information was significantly higher in track B than track A (Figure 4A right). To investigate the cause of the increased spatial information content of the place cells, we examined the out-of-field event rate, the maximum bin event rate, and the field size over the course of the experiment. Both the out-of-field event rate and the field size decreased as the experiment progressed (Supplementary figure 3A, B), while the maximum bin rate did not change significantly (Supplementary figure 3C). For a given recording, we observed a decrease in the proportion of place cells of the population of active cells, as more time was spent in the environment (Figure 4B left). The proportion of active place cells in B_210_ was significantly lower than that of A_200_ (Figure 4B right). This reduction in the proportion of place cells was not accompanied by a concordant reduction in the overall number of active cells in each imaging recording (Figure 4C). The increase in spatial information content of place cells was correlated with the decrease in the proportion of place cells (Figure 4D). This suggests that the encoding power did not change, despite the decrease in the number of active place cells. To examine the combined effect of the changes in spatial information and the proportion of place cells, on the population encoding of space, we trained a maximum likelihood estimator (MLE) for decoding and cross-decoding the neural activity (see the methods for more details). We observed a slight insignificant increase (A_0_ - A_200_: 0.74 cm; B_10_ - B_210_: 3.66 cm) in the decoding error, throughout the experiment (Figure 4E). This suggests that the population of place cells retained its capacity to encode spatial information, despite the substantial reduction in the number of cells. To further assess the effect of the representational drift, we performed a cross-decoding analysis of the place cell population, between recordings (see methods). This revealed more accurate cross-recording decoding in track A than track B (Figure 4F). Taken together, these results suggest that the efficiency of hippocampal representation increases with experience, without compromising its accuracy.

**Figure 4.**
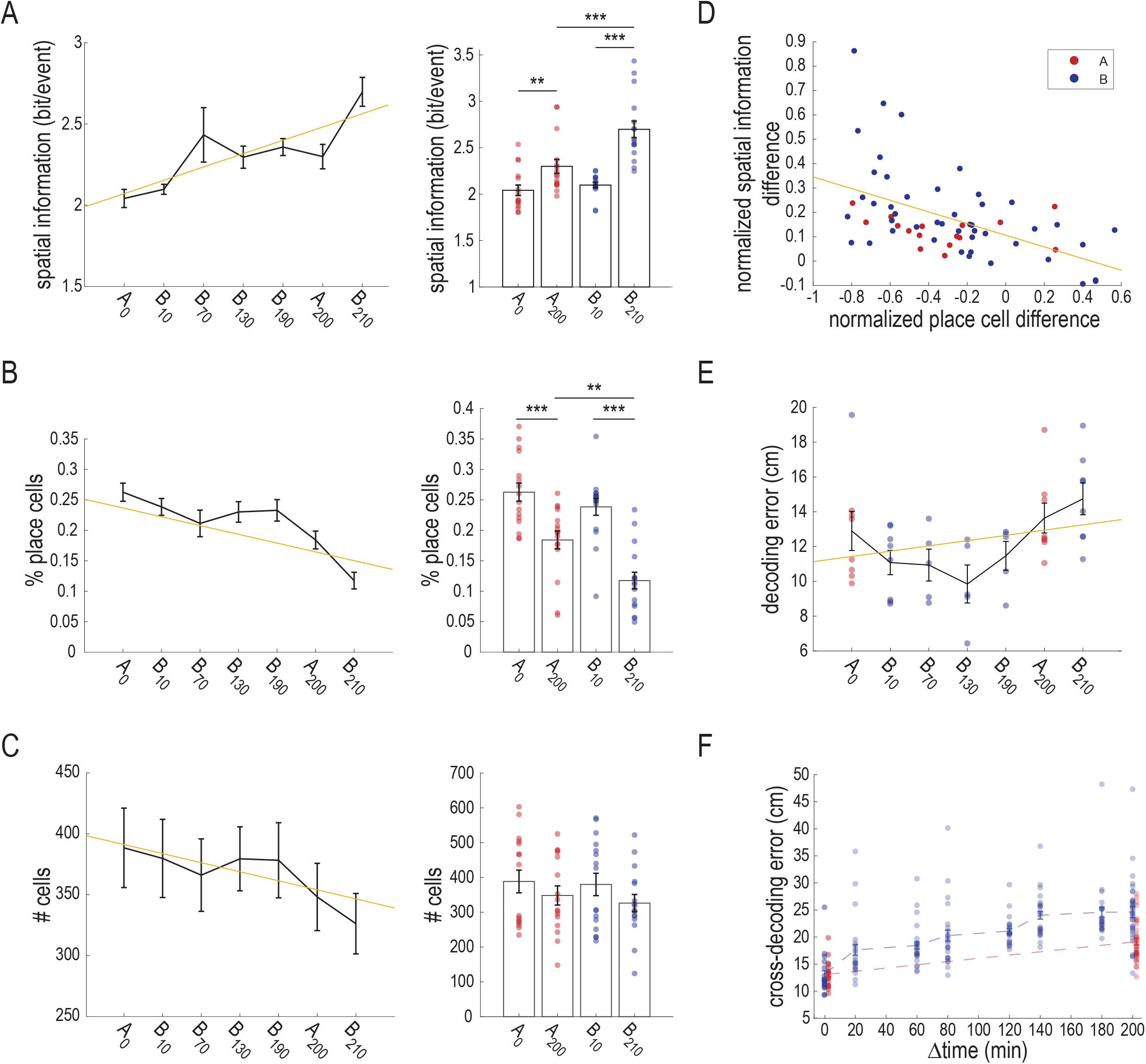
As the time spent in an environment increased, the spatial information content of place cells increased and the number of place cells decreased. A. Spatial information content (n=8 mice) divided into runs north and south throughout the imaging sessions in both contexts. Left: The fitted regression line between spatial information and the time spent in the context was significant (R^2^ =0.23, p<0.001); the spatial information of place cells increased as the time spent in the context increased (β=0.08, p<0.001). Right: bar plot of the spatial information content in the first and last recordings in tracks A and B. The effect of time on spatial information was statistically significant, F(1, 34) =79.29 p<0.001 (two-way repeated measures ANOVA). The spatial information was higher in B_210_ than in B_10_ (mean±s.e.m: B_10_=2.09±0.03; B_210_=2.69±0.08; p<0.001), and in A_200_ than in A_0_ (mean±s.e.m: A_0_=2.04±0.05; A_200_=2.29±0.07; p=0.004). At the end of the experiment, the spatial information was higher in track B than track A (B_210_ - A_200_ p<0.001, T-test with Bonferroni correction for multiple comparisons). B. The proportion of place cells from all the active cells, in the same recording, during all recordings for both contexts. Left: The fitted regression model between the overall proportion of active place cells and the time spent in the context was significant (R^2^=0.293, p<0.001); the proportion of active place cells decreased as more time was spent in the context (β=-0.02, p<0.001). Right: The effect of time on the proportion of place cells was statistically significant, F(1, 34) = 4.80 p=0.035 (two-way repeated measures ANOVA). The proportion of place cells active in each session was smaller in B_210_ than in B_10_ (mean±s.e.m: B_10_=0.23±0.01; B_210_=0.11±0.01; p<0.001), and was also smaller in A_200_ than in A_0_ (mean±s.e.m: A_0_ =0.26±0.01; A_200_ =0.18±0.01; p=0.006). The proportion of place cells in B_210_ was lower than in A_200_ (p=0.009, T-test with Bonferroni correction for multiple comparisons). C. The number of active cells in each session. Left: The regression model for the number of active cells in each recording was not significant (R^2^=0.027, p=0.108). Right: The number of active cells did not differ significantly between recordings, F(1, 34) = 1.62 p=0.212 (two-way repeated measures ANOVA). D. The difference in spatial information content between the first and last recordings in both tracks (n= 62 recordings), normalized by the values of the first recording ([SInfo_last_ – SInfo_first_]/SInfo_first_), plotted as a function of the normalized difference in the proportions of place cells between the first and last sessions in each track ([%PC_last_ – %PC_first_]/%PC_first_), r=-0.515, p<0.001 (Pearson’s correlations). E. Decoding error of the maximum likelihood estimator (MLE) decoder in the imaging sessions. The decoding error increased moderately as more time was spent in the context (R^2^ =0.09, p=0.039), although differences between the groups were not significant (t-test with Bonferroni correction for multiple comparisons). F. Cross-decoding error of the MLE decoder. The cross-decoding error was smaller in track A than track B (p<0.001; t-test with Bonferroni correction for multiple comparisons).

## DISCUSSION

We examined the stability over a number of hours, of spatial representations in the hippocampus, in mice traversing two familiar environments in which they spent substantially different amounts of time. We found that the representations of a track in which the mouse spent relatively little time (20 min. in total, in two visits) were highly correlated between the visits, three hours apart. In contrast, the representations of a track in which the mouse spent a much longer time (about 3 hours), were significantly less correlated between two recordings, three hours apart. This indicates more malleability to change with longer time spent in an environment, even an environment without any noticeable physical difference from a previously experienced one or from the surrounding area.

### Representational drift depends on accumulated experience

Previous studies reported gradual decreases in spatial correlations, with repeated exposures to the same environment^12,14^, over a period of days. In our study we observed a comparable decrease in correlation, but over a period of three hours rather than days. We attribute the steeper decrease in correlations seen in our study to the long time spent in the track. However, while the largest change in representation was in track B, where the mouse spent most of its time, we also observed a change in representation in track A, though it was visited for only short periods. This representational drift may reflect the previously suggested continuous sparsification of the representation, even in the absence of active use, as the brain’s way of compressing experiences, thereby achieving greater efficiency by using less resources^17,18^. The lesser change in representation in track A could also be due to context generalization, as the two tracks were interconnected^19^.

The correlations of the spatial representations gradually decreased as the time spent in a given track increased. This suggests a constant representational drift of the place cell population, as has been shown in consecutive re-exposures to the same environment^13,14^. Moreover, in a similar recording task in a rectangular arena, we found that the decrease in correlation in each half of the arena did not depend on the time spent there. This suggests that the rate of representational drift was related to accumulated experience in the context, rather than to the specific locations within it.

### A decrease in active place cells is accompanied by an increase in spatial information

Previous studies reported that spatial information content increased with repeated exposures to the same environment^20,21^. In our experiment, this effect was also mediated by the time spent in a familiar environment. The increase in spatial information per cell was accompanied by a decrease in the number of overall active place cells. Thus, as time progressed in a specific context, fewer place cells were in use, but their individual information content was higher. We suggest that this increases the efficiency of representations, as more experience is accumulated within an environment, thus reducing the resources needed as the network becomes more tuned.

To test this efficiency hypothesis, we used a maximum-likelihood decoder to decode the animals’ position at each time point. We found that the decoding quality did not substantially degrade between the start and the end of the experiment, suggesting that the increase in spatial information per cell compensated for the decrease in the number of place cells. This hints to the possibility that the hippocampus balances the amount of spatial information within its network by decreasing the number of place cells with accumulated experience, while increasing the spatial information per cell, such that the total information does not change much, but the load on the network is reduced^21,22^.

In summary, our results indicate that even in the absence of perceived change, memory of a given environment is constantly updated, thereby inducing a continuous representational drift that is dependent on the amount of time spent in that environment. This resonates well with the phenomenon of lability to change after activation, known as reconsolidation^23^. The notion that hippocampal memory may be subject to change, when active, indicates a tight link between activity within context, and plasticity of hippocampal representations.

## STAR methods

Contact for Reagent and Resource Sharing: Further information and requests for resources, reagents and Matlab code should be directed to and will be fulfilled by the Lead Contact.

## METHODS

### Mice and surgical procedures

All the surgical and experimental procedures were approved by the Animal Care and Use Committee of the Technion – Israel Institute of Technology. All the mice were from the same C57BL/6 background from Jackson Laboratories. The mice were aged 8-12 weeks at the start of the procedures. They were housed separately and underwent two surgical procedures under isoflurane anesthesia (1.5-2% volume) accompanied by buprenorphine analgesia (0.7 mL of 1:60 saline-diluted 30 mg/ml buprenorphine).

The mice were injected with the viral vector AAV1-syn-jGCaMP7f-WPRE (~1^13^ vg/mL, Addgene) into the dCA1 (stereotactic coordinates: −2.1 mm anteroposterior, 1.25 mm mediolateral, −1.4 mm dorsoventral from bregma) of the hippocampus using a pulled glass micropipette. These injections were 250-300 nL in volume, at a rate of 0.05 μl a minute. Two weeks after the injections, the mice underwent a second surgery. A craniotomy (1mm in diameter) was performed in the same coordinates as the GCaMP injection. We removed the dura and cortex above the CA1 by suction with a 29-gauge blunt needle while constantly washing the exposed tissue with sterile PBS. Then we implanted a GRIN lens (ProViewTM Lens Probes 1.0mm diameter, ~4.0mm length, Inscopix, Palo Alto, CA) directly above the CA1, and sealed the space between the skull and the lens with kwik-sill (WPI, Sarasota, FL). Afterwards, we used Metabond (Parkell, Edgewood, NY) to cover the exposed skull and the lateral sides of the lens. Two weeks after, a baseplate (Inscopix) was installed above the lens. This was done by lowering the miniature microscope (Inscopix) until it reached an in-focus imaging plane, after which the baseplate was attached to the Metabond (Parkell) covered skull using light-cured dental cement.

### Food restriction, training, and reward habituation

After at least one-week recovery following the implantation surgery, the animals were food restricted to 2.5-3 g of food pellets (Altromin 1324 IRR complete animal feed for laboratory animals) to maintain 85%-90% of their original body weight. The mice were trained to run back and forth in the linear track, receiving small banana-flavored food pellet rewards at both ends of the track. This was done for 3-5 days prior to the experiment to familiarize the animals with the track and to obtain good coverage during the experiments. Training was finished when the animals could do at least 10 runs back and forth (20 meters) in 10 minutes while eating all the food rewards offered. After completing the training, the animals were ready for the experimental phase.

### Experimental setup

The experimental setup consisted of a custom-made maze composed of two linear tracks measuring 100 cm in length, 10 cm in width, and 10 cm in height. The tracks were painted black and suspended 40 cm above the ground using a small table. We used overhead lights to illuminate the track, and black curtains surrounded the track from all sides. At the beginning of each experiment, the animals were first connected to the miniature microscope (Inscopix) in the home cage while we made sure to return to the same field of view before the start of the imaging recording. Mice were placed inside the intermediate chamber, and then released into track A where they spent 10 minutes, while the activity of their cells were imaged (A_0_). We then opened the doors to track B, which the mice traversed for 200 minutes. The activity of cells in track B was imaged for the first 10 minutes only (B_10_). After 200 minutes, the mice returned to track A through the intermediate chamber, for 10 additional minutes of imaging (A_200_), and finally to track B, where the cells were imaged for an additional 10 minutes (B_210_), resulting in a total of four 10-minute recordings (Figure 1B). The linear track surface was cleaned after each session with 70% ethanol. For the 2D experiments, the same behavioral protocol described above was repeated, only in a different maze. This maze consisted of two similar rectangular arenas connected by a door (Figure 3A).

### Ca^2+^ imaging and processing of the data

We imaged the calcium signal using a miniature microscope (nVista, Inscopix) at 20 Hz. Recordings were synchronized with the behavioral camera mounted above the track. We processed the imaging data using the Inscopix data processing software (IDPS) (1.3.1) and custom written MATLAB codes. To ascertain similar processing for all recording epochs, analysis of imaging data was combined for all recording epochs of the same experiment. For processing the imaging data, we followed previously described routines^13^. Specifically, we spatially downsampled the videos in each dimension. Then we applied a 3×3 median filter to fix any defective pixels due to unequal illumination or defects in the sensor itself. Subsequently, a spatial bandpass filter was applied using the IDPS (low cut-off: 0.005, high cut-off: 0.5 *pixel*^−1^) to achieve a smoothed version of the original video. This enhanced the appearance of the blood vessels and was later used to make the motion correction more effective. We then applied a motion correction algorithm using IDPS (correction type: translation and rotation; reference region with maximum registration value (r = 20 pixels)). For calcium signal extraction from putative single CA1 neurons, we used the constrained non-negative matrix factorization – extended algorithm (CNMF-E) using MATLAB^24,25^. The algorithm isolated the putative single units from the processed imaging videos automatically. Isolated putative units that did not match spatial or temporal features of the neurons were discarded and not used in subsequent analyses. All the analyses used the deconvolved activity inferred by CNMF-E.

### Place fields

We computed spatial firing rates (rate maps) by dividing the one-dimensional track to 25 spatial bins (4 cm/bin). We divided the neural and behavioral data by conducting direction runs to the left and to the right. Events that occurred and position information that accumulated during non-movement epochs (< 1 cm/sec) were excluded. We also excluded the last bins at the two ends of the tracks, where the mice were generally stationary due to consumption of food rewards. We then divided the number of events in each spatial bin by the amount of time the mouse spent in that bin per direction. We used a Gaussian smoothing factor (sigma = 3 cm) for each bin and normalized each place field by its maximum value. Unvisited spatial bins were marked for exclusion in later analyses. Place fields with less than five events (Ca^2+^ imaging) were excluded from the analysis. For place cells included in the analysis, we calculated the spatial information (SI, bits/event) for each cell, as previously described:

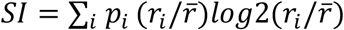

Where *r_i_* is the rate of the neuron in the ith bin; *p_i_* is the probability of the mouse being in the ith bin (time spent in the *i*-th bin/total session time); 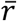 is the overall mean rate; and i indicates running over all the bins. We then performed a temporal shuffling procedure for each rate map, to test for statistical significance of spatial information. Event timestamps were moved in a random non-repeating circular shift relative to the position time in each trial, 1000 times for each cell. We computed rate maps and spatial information for every iteration. A cell was considered as spatially modulated during a trial if its spatial information score was higher than 950 of the shuffled data instances of spatial information, for a significance level of p < 0.05.

### Single-cell correlation

We calculated the correlation between the activity of each two corresponding neurons in a particular imaging recording, using the rate maps of these neurons. We calculated the Spearman’s correlation between the maps for each two neurons. Then we batched all the cell-pair correlations of all the mice from the same sessions together.

### Positional shift

For each neuron, we calculated the positional shift as the absolute difference between the positions of the peak event rate (Gaussian smoothing factor sigma = 3 cm) on the track in two recordings. Then we averaged all the positional shifts for the neurons of all the mice in the same sessions.

### Decoding position from activity

To decode the position of the mouse from the neural activity, we used a maximum likelihood estimation (MLE) decoder. We assumed that neural events were uncorrelated, which is practically untrue and diminishes decoding accuracy but greatly simplifies calculation time. In addition, because events are so sparse that most time bins are either empty of activity or contain only one event, we used a maximum filter (*size* = 250[*ms*]) over each neuron’s activity vector. The likelihood function can then be written as:

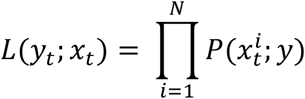

Where 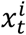 is a binary variable that represents whether neuron *i* fired at time *t*, *N* is the number of place cells used for the decoding, 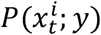 is the *y* bin of the *i_th_* neuron normalized unsmoothed rate map if the neuron fired, and the inverse normalized rate map if it did not fire. The position can then be decoded using maximum likelihood.

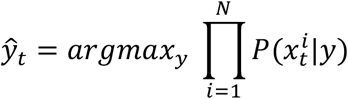

Because the probabilities are very small, we used the following adjusted formula to avoid numerical errors; and as the transformation is monotonous, the decoding is not changed:

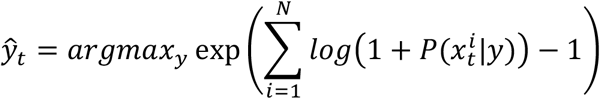

To test the decoder’s error, we split the recording data such that the rate maps were estimated based on 75% of the linear track traversals (in each direction) and the MSE was calculated over the remaining 25%. For the cross decoder, we calculated the rate maps based on the first recording and the error over the second recording.

## Acknowledgements

We thank Yaniv Ziv for reading and commenting on the manuscript. This research was supported by the ISRAEL SCIENCE FOUNDATION (grants Nos. 2655/18 and 2183/21 to DD, and 1442/21to OB), by the German-Israeli Foundation (GIF I-1477-421.13/2018) to DD, by a grant from the US-Israel Binational Science Foundation (NIMH-BSF CRCNS BSF:2019807, NIMH:R01 MH125544-01 to DD), by an HFSP research grant (RGP0017/2021) to OB, A Rappaport Institute Collaborative research grant to DD, by Israel PBC-VATAT and by the Technion Center for Machine Learning and Intelligent Systems (MLIS) to DD and OB, by the Prince Center for the Aging Brain, and by a University of Michigan – Israel Partnership for Research and Education Collaborative Research stipend to DK.

---

**Supplementary figure 1:**
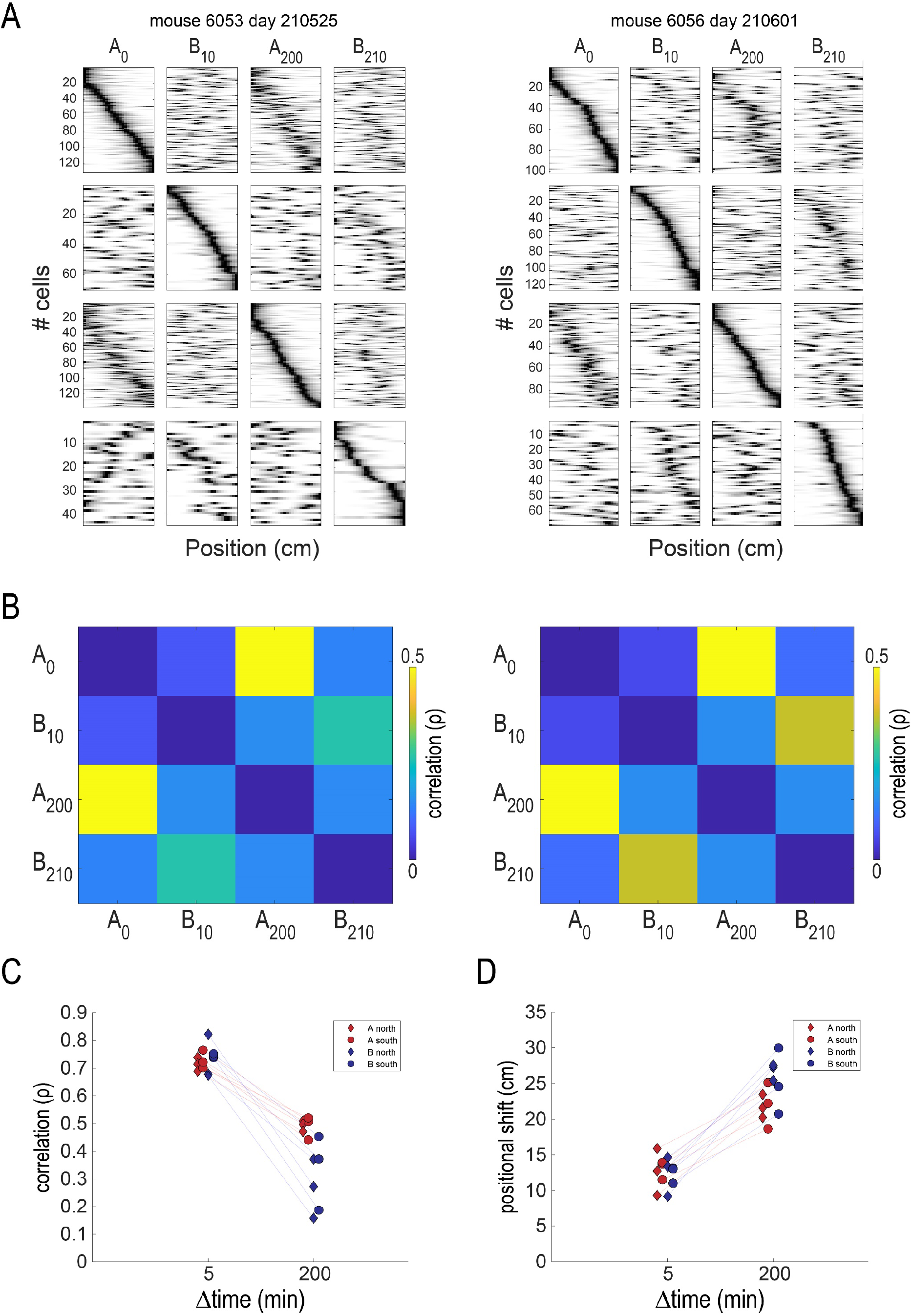
Rate maps and correlation examples (related to figure 1). A. *Two examples of rate maps across recordings of two mice. In each row are shown average rate maps for individual cells along the linear track of place cells that were active during that specific session, normalized by peak activity*. B. *Pairwise correlations of the individual cells between recordings, for the same mice shown in a*. C. *Pairwise single-cell correlations, according to mice and to runs in different directions (north and south)*. D. *Positional shift of the center of mass of all the place cells, according to mice and to runs in different directions (north and south)*.

**Supplementary figure 2:**
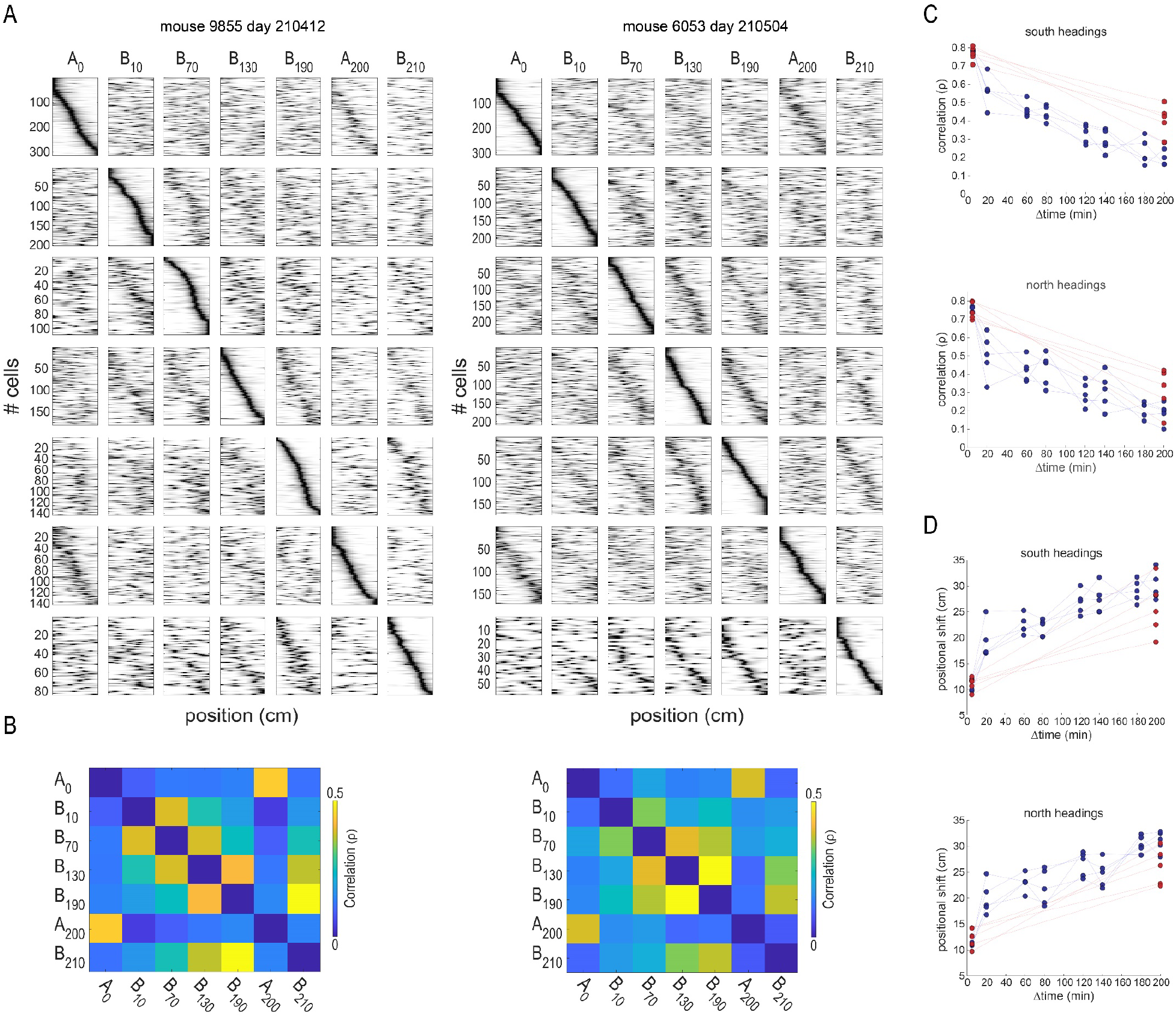
Examples of rate maps and correlations of the extended protocol (related to figure 2) A. *Two examples of rate maps across recordings of two mice in the extended imaging protocol. In each row are shown average rate maps for individual cells along the linear track of place cells that were active during that specific recording, normalized by peak activity*. B. *Pairwise correlations of individual cells between different imaging sessions for the same mice shown in a*. *C. Pairwise single-cell correlations*, as a function of the time difference between sessions, for mice in the south-heading runs (top) and the north-heading runs (bottom). *Positional shift of the center of mass of all the place cells*, for mice in the south-heading runs (top) and the north-heading runs (bottom).

**Supplementary figure 3:**
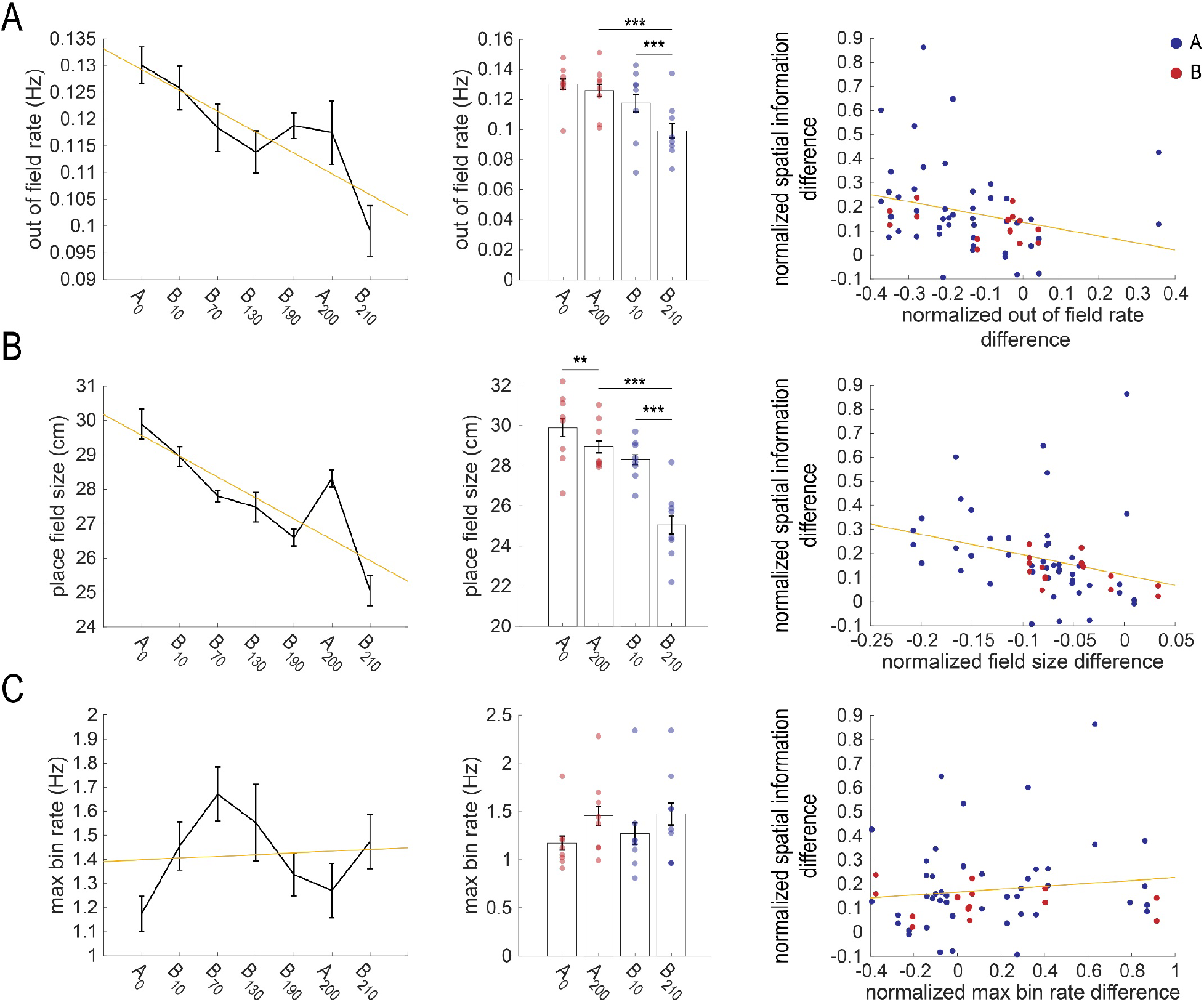
The effect of place field size, maximum firing rate of each bin, and out of field firing on spatial information content of place cells (related to figure 4) A. *The out-of-field event rate (n=8 mice) according to runs north and south along the recordings in both contexts. Left: The fitted regression line between the out-of-field event rate and the time spent in the context was significant (R^2^ =0.196, p<0.001). The out-of-field event rate of place cells decreased as the time spent in the context increased (β=-0.004, p<0.001). Middle: bar plot of the out-of-field event rate in the first and last recordings in tracks A and B. The effect of time on the out-of-field event rate was statistically significant, F(1, 30) = 22.5 p<0.001 (two-way repeated measures ANOVA). The out-of-field event rate was lower in* B_210_ *than in B_10_ (mean±s.e.m: B_10_=0.12±0.004; B_210_=0.09±0.004; p<0.001), but not in A_200_ compared to* A_0_ (*mean±s.e.m: A_0_=0.13±0.003; A_200_=0.11±0.005; p=0.237). At the end of the experiment, the spatial information was greater in track B than track A* (*B*_210_ - A_200_ *p<0.001, T-test with Bonferroni correction for multiple comparisons*). Right: The difference in *spatial information content between the first and last sessions in both tracks, normalized by the values of the first recording (see figure 4d for the equation), plotted as a function of the normalized difference in the out-of-field event rate between the first and last recordings in each track. r=-0.26, p=0.037 (Pearson’s correlations*). B. *The max bin rate for runs north and south along the imaging sessions in both contexts. Left: the fitted regression line between the max bin rate and the time spent in the context was not significant (R^2^=0.007, p=0.58). Middle: bar plot of the max bin rate in the first and last imaging sessions in tracks A and B. The effect of time on the max bin rate was not statistically significant, F(1, 30) = 0.576 p=0.454 (two-way repeated measures ANOVA)*. Right: *the difference in spatial information content between the first and last recordings in the two tracks, normalized by the values of the first recording plotted as a function of the normalized difference in the max bin rate between the first and last recordings in each track. r=0.13, p=0.312 (Pearson’s correlations)*. C. *Field size for the runs north and south along the imaging sessions in both contexts. Left: The fitted regression line between field size and the time spent in the context was significant (R^2^=0.403, p<0.001). The out-of-field event rate of place cells decreased as the time spent in the context increased (β=-0.607, p<0.001). Middle: bar plot of the field size in the first and last recordings in tracks A and B. The effect of time on the out-of-field event rate was statistically significant, F(1, 30) = 103.87, p<0.001 (two-way repeated measures ANOVA). The field size was lower in* B_210_ *than in B_10_ (mean±s.e.m: B_10_=28.9±0.3; B_210_=25.05±0.43; p<0.001) and in* A_200_ *than in* A_0_ (*mean±s.e.m: A_0_=29.9±0.44; A_200_=28.3±0.25; p=0.001). At the end of the experiment, the spatial information was greater in track B than track A* (*B*_210_ - A_200_ *p<0.001, T-test with Bonferroni correction for multiple comparisons*). Right: *The spatial information content difference between the first and last recordings in both tracks, normalized by the values of the first session (see figure 4d for equation), plotted as a function of the normalized difference in the out-of-field event rate between the first and last recordings in each track. r=-0.288, p=0.023 (Pearson’s correlations)*.

## Notes

### Competing Interest Statement

The authors have declared no competing interest.

